# Sanguinarine activates ATM/ATR-mediated CHK-1 signaling to drive p53-dependent apoptosis in the *C. elegans* germline

**DOI:** 10.64898/2026.01.26.701699

**Authors:** Raghad El Ghali, Mahmoud Izadi, Zainalabedin Alrayyes, Tayyiba Akbar Ali, Shahab Uddin, Ehsan Pourkarimi

**Affiliations:** Division of Genomics and Translational Medicine, College of Health and Life Sciences, Hamad Bin Khalifa University, Qatar Foundation, Doha, Qatar; Translational Research Institute, Academic Health System, Hamad Medical Corporation, Doha, Qatar

**Keywords:** Sanguinarine, DNA damage, apoptosis, cancer, *Caenorhabditis elegans*

## Abstract

Sanguinarine (SNG) is a natural component belonging to the benzophenanthridine alkaloids. In recent years, due to its remarkable biological activities, it has gained wide interest in the pharmaceutical industry. Various studies have reported its potential as a therapeutic agent in treating chronic human diseases such as cancer. SNG is widely reported to cause programmed cell death in various cancer cell lines. The mechanism by which SNG triggers apoptosis remains poorly elucidated, especially *in vivo*. Previous studies reported that sanguinarine induces apoptosis by increasing reactive oxygen species (ROS). In this study, we aimed to characterize the effects of SNG using an *in vivo Caenorhabditis elegans* (*C. elegans*) model. Treating *C. elegans* with various SNG concentrations resulted in apoptotic cell death in the proliferative germline. Interestingly, SNG-induced apoptosis depends on the core apoptotic machinery initiated by the DNA-damage-induced activity of the p53/CEP-1 protein. We have also demonstrated that the increase in germ cell apoptosis is caused by elevated levels of reactive oxygen species (ROS) following SNG treatment. Notably, the apoptotic phenotype induced by SNG was resolved upon treatment with the ROS scavenger. Altogether, our study demonstrates that SNG increases ROS, leading to activation of DNA damage-induced apoptosis in the proliferative germline of *C. elegans*.

## Introduction

Sanguinarine (SNG) is a benzophenanthridine alkaloid originally isolated from *Sanguinaria canadensis* [1-6]. More recently, SNG has attracted substantial interest due to its reported antitumor activity. Numerous *in vitro* studies have demonstrated that SNG inhibits tumor growth and induces apoptosis in a wide range of cancer cell lines, including lung adenocarcinoma, colon, skin, prostate, and basal-like breast cancers [7-9]. SNG was shown to cause cell cycle arrest and induce apoptotic death in human prostate carcinoma cells [10]. Consistently, SNG has also been shown to inhibit tumor growth in mouse xenograft models [11]. However, there are conflicting reports regarding the specificity of SNG against cancer cells and the extent of its general cytotoxicity in non-cancerous tissues [12-14]. Moreover, adverse effects such as embryonic and developmental toxicity, as well as increased apoptosis, were described in recent studies in zebrafish and mouse [13, 15, 16].

Over the past years, the antineoplastic effect of SNG has been linked to multiple signaling pathways, including inhibition of PI3K/AKT, MAPK, NF-κB, and JAK/STAT signaling *in vitro* [16-21]. SNG has also been reported to induce cell cycle arrest at multiple stages, including G1/S and G2/M arrest in prostate cancer cells and G0 arrest in breast cancer cells [10, 22]. Moreover, SNG appears to have a dose-dependent cell death effect, as treatment of cancer cells at low concentrations (1-4 μM) induces caspase-dependent apoptosis, while higher concentrations (9-17 μM) have been associated with necrotic cell death [23]. Although SNG-induced apoptosis has largely been reported as p53-independent, p53 dependency has been observed in specific contexts. Notably, in prostate cancer LNCaP cells, SNG induces p53-dependent cell death, underscoring the need for further investigation in genetically intact systems [24]. The cytotoxic effects of SNG are attributed largely to its ability to elevate reactive oxygen species (ROS).

ROS are physiologically important in various cellular processes, notably in cellular proliferation and survival; however, excessive ROS levels have adverse effects and can lead to oxidative damage of cellular components, inducing DNA lesions and apoptosis [25]. Cancer cells often have elevated levels of ROS due to their high metabolic demand; therefore, ROS-inducing therapies, such as sanguinarine, push the levels of ROS in cancer cells beyond their tolerance threshold, leading to apoptosis [26]. Despite extensive investigation, the precise relationship between SNG exposure, DNA damage signaling, and checkpoint activation remains poorly defined. Most mechanistic studies have been conducted in transformed cancer cell lines that frequently harbor defective DNA damage checkpoints and pre-existing genomic instability, complicating the interpretation of whether SNG directly engages canonical DNA damage pathways or triggers apoptosis through checkpoint-independent stress responses. We therefore examined the pro-apoptotic effects of SNG in an intact organismal setting using the *Caenorhabditis elegans* (*C. elegans*) germline. Our data shows that SNG activates the canonical DNA damage checkpoint pathway through the upstream kinases ataxia telangiectasia mutated (ATM) and ataxia telangiectasia and Rad3-related (ATR), resulting in CHK-1 activation and subsequent CEP-1/p53-dependent, caspase-mediated apoptosis. Notably, SNG treatment selectively increases reactive oxygen species in proliferative germ cells, rather than inducing a global oxidative stress response. All in all, these findings establish that SNG triggers DNA damage-induced apoptosis through checkpoint-dependent mechanisms and reveal a functional interplay between ROS and genome surveillance pathways in shaping cellular responses to this widely studied natural compound.

## Results

### Induction of caspase-dependent apoptosis by SNG in *C. elegans* germline

To investigate if sanguinarine induces apoptosis in the proliferating germline of *C. elegans*, we treated the *ced-1::gfp s*train with various concentrations of SNG, ranging from 8 μM to 25 μM SNG, with DMSO as a control. CED-1 (cell death abnormality protein 1) is a transmembrane protein essential for apoptotic engulfment that accumulates around the dying cell, making it a well-established cytological marker to identify apoptotic cells [27, 28]. Notably, treatment with 20 μM and 25 μM for 24 hours resulted in a robust and significant increase in germ cell apoptosis compared to control (p < 0.0001) (**Figure 1a**). Notably, exposure to 25 μM SNG did not cause germline disorganization or any changes in the gross anatomy; therefore, SNG concentration of 25 μM was chosen as the reference dosage for further experiments (**Figure 1b**).

**Figure 1.**
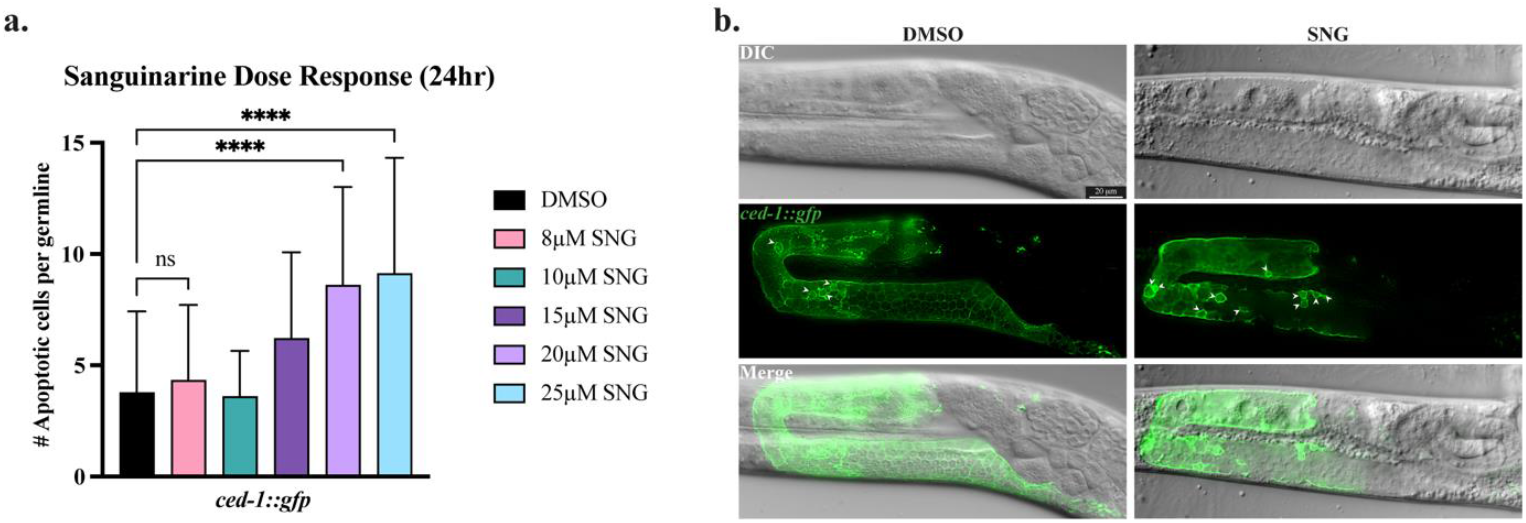
Sanguinarine treatment induces apoptosis in the germline. **a**. Number of apoptotic cells per germline in DMSO and SNG-treated *ced-1::gfp* worms. SNG concentrations of 20 μM and 25 μM show a significant increase in apoptotic cell count (**** p<0.0001). **b**. Microscopic images of *ced-1::gfp* germlines of worms treated with DMSO or sanguinarine (25 μM). White arrowheads indicate the apoptotic cells in the pachytene region.

In adult *C. elegans*, the germline is the only proliferative tissue and is spatially organized along a distal–proximal axis, with mitotically dividing cells at the distal end and meiotic progression toward the proximal gonad [29]. Importantly, germ cell apoptosis is restricted to the pachytene region of meiotic prophase I of the germline [28]. In *C. elegans*, apoptotic cells are executed through the conserved core apoptotic machinery composed of CED-9/Bcl2, CED-4/Apaf1 and the most downstream protein, the sole *C. elegans* caspase, CED-3 [28, 30, 31]. To test if the SNG-induced germ cell apoptosis requires CED-3/caspase activity, we treated the *ced-1::gfp;ced-3*(*n2452*) strain with 25 μM SNG. *ced-3*(*n2452*) is a *ced-3* null allele that abolishes all programmed cell death. As expected, loss of *ced-3* completely suppressed SNG-induced apoptosis, indicating that the apoptotic response to SNG is caspase-dependent (p < 0.0001) (**Fig 2a, 2b**).

**Figure 2.**
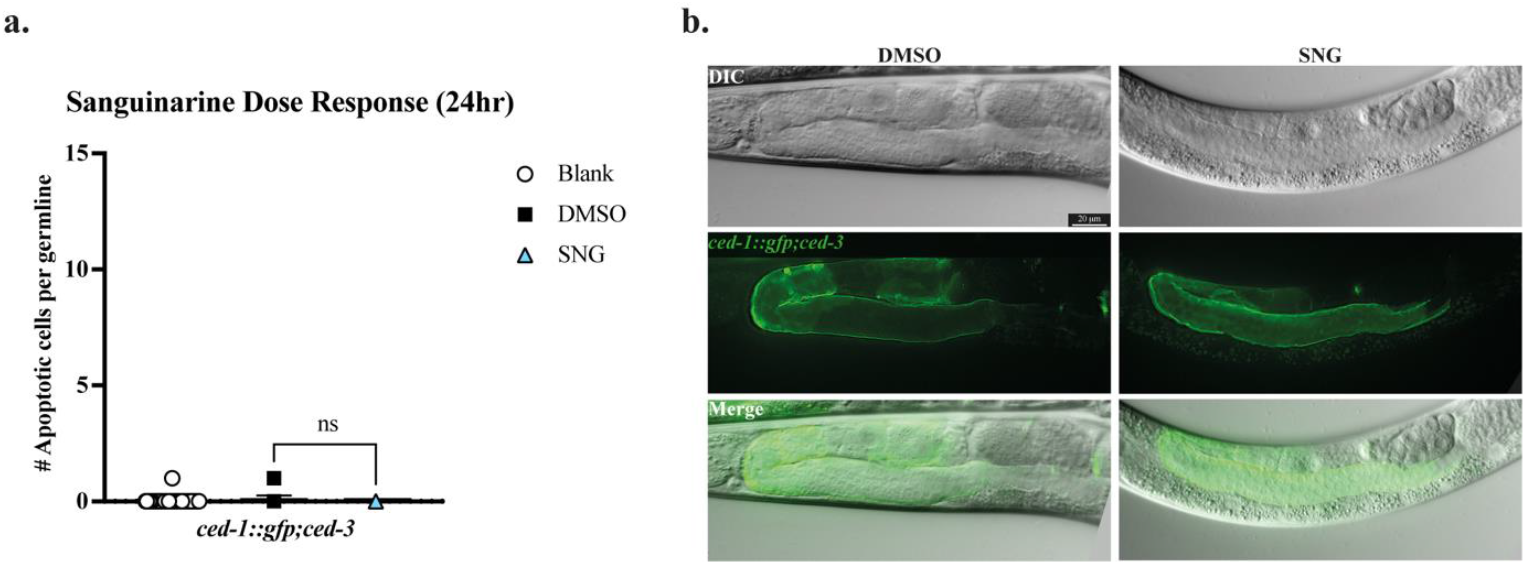
Sanguinarine-induced apoptosis is caspase-dependent. **a**. No significant difference in apoptotic cell counts between SNG-treated worms and DMSO control in a caspase-null background was observed. **b**. Microscopic images of caspase-null *ced-1::gfp;ced-3* germlines upon DMSO and SNG treatment.

### SNG does not induce developmental apoptosis in *C. elegans*

During the invariant development of *C. elegans*, 131 somatic cells undergo programmed cell death as part of the normal developmental program [32, 33]. This developmental apoptosis depends on the core apoptotic machinery [34]. To test whether SNG-induced apoptosis is restricted to the proliferative germline or also affects developmental apoptosis, we measured developmental apoptosis upon SNG exposure. To this end, we used engulfment-defective *ced-1* (*e1735*) mutants, in which apoptotic cells persist due to impaired engulfment and clearance, allowing reliable quantification of developmental cell death [27]. Embryos were collected from the SNG-treated hermaphrodites, and the developmental apoptotic cells of the pharyngeal region were measured in the first larval stage (L1). Interestingly, no significant difference was observed in pharyngeal apoptotic counts between the L1 larvae of sanguinarine-treated worms and those of untreated or DMSO-treated controls (**Supplemental Figure S1**). These results indicate that SNG selectively induces apoptosis in the proliferating germline without affecting developmental apoptosis in somatic tissues.

### CEP-1/p53 is required for sanguinarine-induced germline apoptosis

Previous studies in cancer cell lines have demonstrated that sanguinarine-induced apoptosis is largely independent of p53, although conflicting observations have been reported [24]. To test if SNG-induced germline apoptosis depends on CEP-1/p53 activity, apoptotic corpses were measured in the *cep-1*/p53-null carrying *ced-1::gfp* strain, *ced-1::gfp;cep-1*. Interestingly, loss of *cep-1*/p53 resulted in a significant reduction in SNG-induced apoptosis compared to SNG-treated *ced-1::gfp* controls (p<0.0001) (**Figure 3a, 3b**). These results demonstrate that, in contrast to many cancer cell line data, sanguinarine-induced apoptosis in the *C. elegans* germline is dependent on the activity of CEP-1/p53.

**Figure 3.**
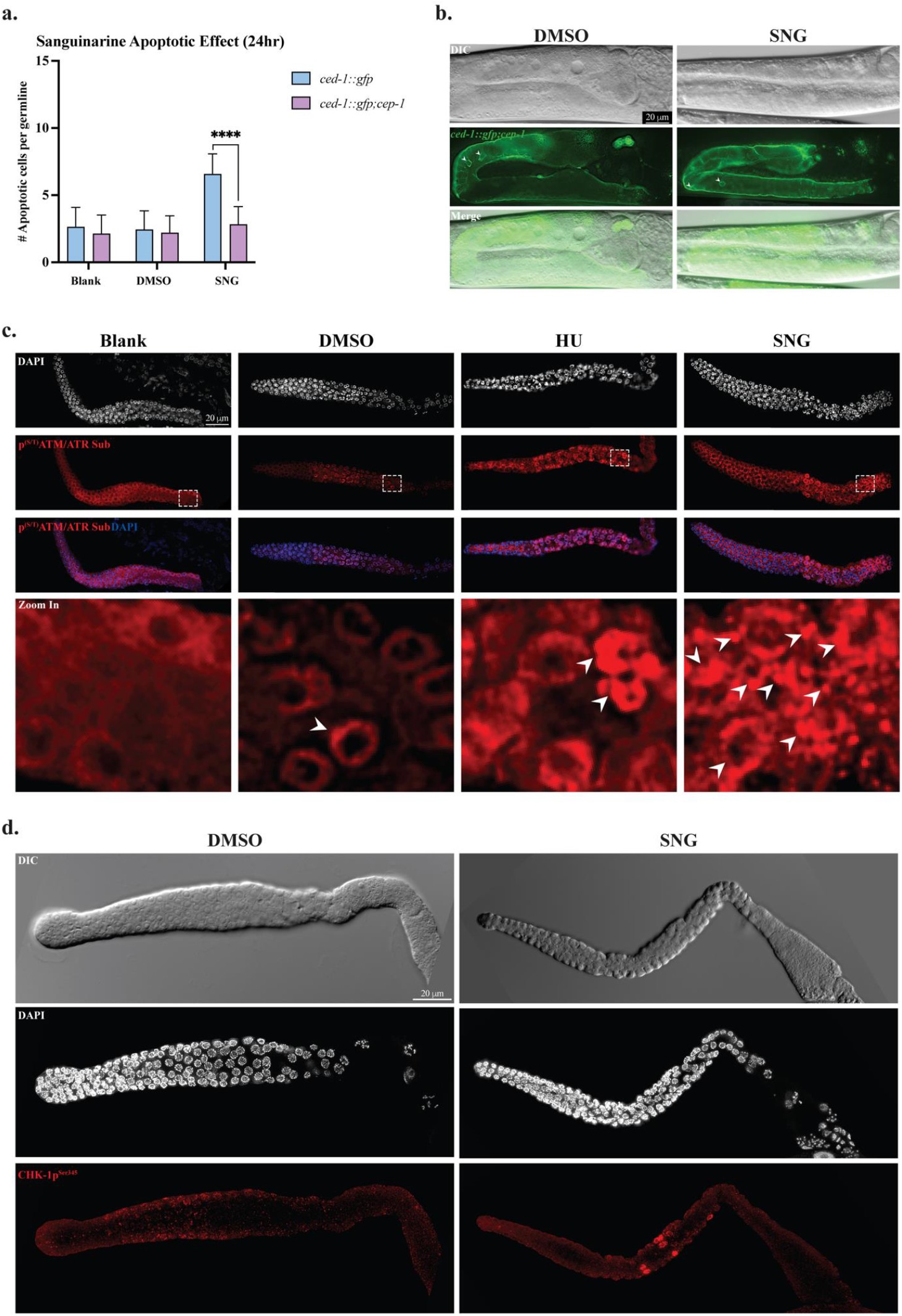
Sanguinarine Treatment Induces DNA Damage Checkpoint Activation through ATM and ATR Signaling. **a**. Significant decrease in germline apoptosis in the *cep-1*/p53-null background upon SNG-treated worms (**** p < 0.0001). **b**. Representative microscopic images of the germlines of *cep-1*/p53-null *ced-1::gfp;cep-1* worms treated with SNG or DMSO. **c**. Representative immunofluorescence images of adult *C. elegans* germlines treated with Blank, DMSO, 25 mM hydroxyurea (HU), or 25 µM sanguinarine (SNG). Germlines were stained for the phosphorylated ATM/ATR substrate motif ((S/T) Q) (red) and counterstained with DAPI (blue). Robust phospho-ATM/ATR signals were observed in HU-and SNG-treated germlines, whereas controls showed minimal signal. Lower panels show zoom in the pachytene region (shown in dashed box) and merged images. **d**. Immunostaining of dissected germlines from DMSO- and SNG-treated worms showing increased CHK-1p^Ser345^ phosphorylation upon SNG treatment. Scale bars, 20 µm.

### Sanguinarine treatment induces the DNA damage checkpoint pathway through activation of ATM and ATR protein kinases

Activation of p53 typically occurs by the activity of the evolutionarily conserved DNA damage response (DDR) pathway mediated by the upstream sensor kinases, ATM and ATR [35]. Given that SNG induces CEP-1/p53–dependent germline apoptosis, we hypothesized that SNG activates the canonical DDR pathway in *C. elegans*. Upon genotoxic stress, ATM and ATR phosphorylate multiple downstream substrates, including p53 and DNA checkpoint proteins, at conserved serine/threonine followed by glutamine ((S/T)Q) motifs [36, 37]. To directly test whether SNG activates ATM/ATR signaling, we assessed phosphorylation of ATM/ATR substrates using an antibody that recognizes the conserved phospho-(S/T)Q motif [37]. Immunostaining was performed on dissected germlines from SNG-treated worms and compared to untreated or DMSO-treated controls. As a positive control, worms were exposed to 25 mM hydroxyurea (HU), a well-established genotoxic agent that robustly activates ATM/ATR signaling (**Figure 3c**).

Among the key downstream targets of ATM/ATR is Checkpoint Kinase 1 (CHK-1), which is phosphorylated at serine 345 in response to DNA damage. This modification is required for checkpoint activation and subsequent CEP-1/p53–dependent apoptosis. Consistent with activation of the canonical checkpoint cascade, we observed robust CHK-1 phosphorylation following SNG treatment [38-40]. To test whether SNG treatment activated the canonical DNA damage checkpoint pathway, we examined CHK-1 phosphorylation. Germlines from SNG-treated animals were immunostained with an antibody against human CHK-1 phosphorylated at serine 345 (CHK-1p^Ser345^), which cross-reacts with the corresponding residue in *C. elegans* CHK-1 [40-42]. SNG treatment resulted in a robust CHK-1p^Ser345^ signal, whereas no detectable phosphorylation was observed in untreated or DMSO-treated controls (**Figure 3d**). These results indicate that SNG activates the DNA damage checkpoint pathway by the activity of ATM/ATR kinases and provide mechanistic support for the observed CEP-1/p53–dependent germline apoptosis.

### Increased ROS production in the germline in response to sanguinarine treatment

Sanguinarine has been reported to induce ROS production in human cell lines and in *C. elegans* at concentrations as low as 0.2 μM [12, 43]. To test if SNG-induced apoptosis is mediated by elevated ROS, we assessed ROS levels in SNG-treated worms using CellROX™ Green, a cell-permeant dye that exhibits bright fluorescent light upon oxidation by ROS. Interestingly, quantification of the whole-worm fluorescence revealed no significant difference between the untreated and SNG-treated worms and only a modest increase relative to DMSO-treated controls (**Figure 4a, 4b**). In contrast, and to our surprise, analyzing the germline revealed a striking and significant difference in ROS levels in SNG-treated animals compared to both untreated and DMSO-treated controls (p<0.001, p<0.0001, respectively) (**Figure 4c, 4d, 4e**). This finding indicates that SNG selectively induces ROS production in the germline rather than the somatic cells.

**Figure 4.**
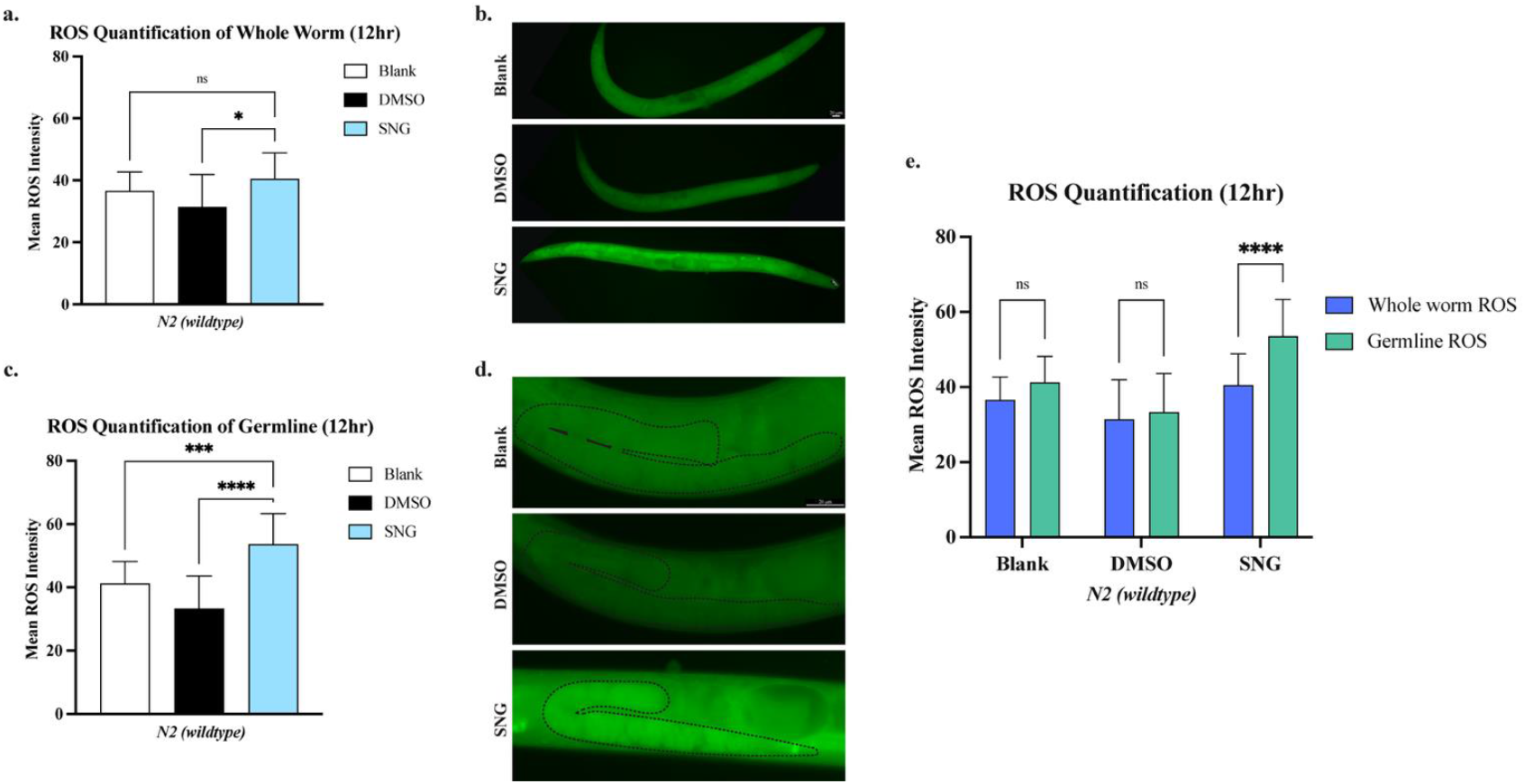
Increase in ROS level upon sanguinarine treatment. **a**. ROS measurement in the whole worms. **b**. Whole worms stained with CellROX Green for ROS detection upon 12h SNG treatment. **c**. The intensity of ROS measured in germline shows a significant increase in SNG-treated worms compared to Blank and DMSO controls (*** p<0.001, and **** p<0.0001, respectively). **d**. Representative images of CellROX Green signal intensity in the germline **e**. Mean ROS intensity is significantly higher in the germline of SNG-treated worms compared to the whole worm (**** p<0.0001).

Because the *C. elegans* germline is the only proliferative tissue in an adult worm, we next asked if mitotic activity is required for SNG-induced ROS generation. To test this, we pretreated worms with 5-fluorouracil (5-FU) to block germ cell mitosis prior to SNG treatment. Notably, inhibition of mitotic proliferation did not suppress SNG-induced ROS accumulation in the germline (**Supplemental Figure S2**), indicating that ROS induction by SNG occurs independently of ongoing cell division.

### ROS scavenger suppresses sanguinarine-induced germline apoptosis

Given the marked increase in germline-specific ROS following SNG treatment, we asked whether oxidative stress contributes to SNG-induced germline apoptosis. To address this, we first examined whether scavenging ROS could suppress SNG-induced oxidative stress in the germline. Wild-type animals were treated with the antioxidant N-acetyl-L-cysteine (NAC), and ROS levels were quantified in the presence or absence of SNG. As expected, NAC treatment resulted in a robust and significant reduction in SNG-induced germline ROS compared to SNG treatment alone (p<0.0001; **Figure 5**). Moreover, NAC suppressed SNG-induced oxidative stress in a dose-dependent manner, with increasing NAC concentrations progressively reducing germline ROS levels (**Figure 5**).

**Figure 5.**
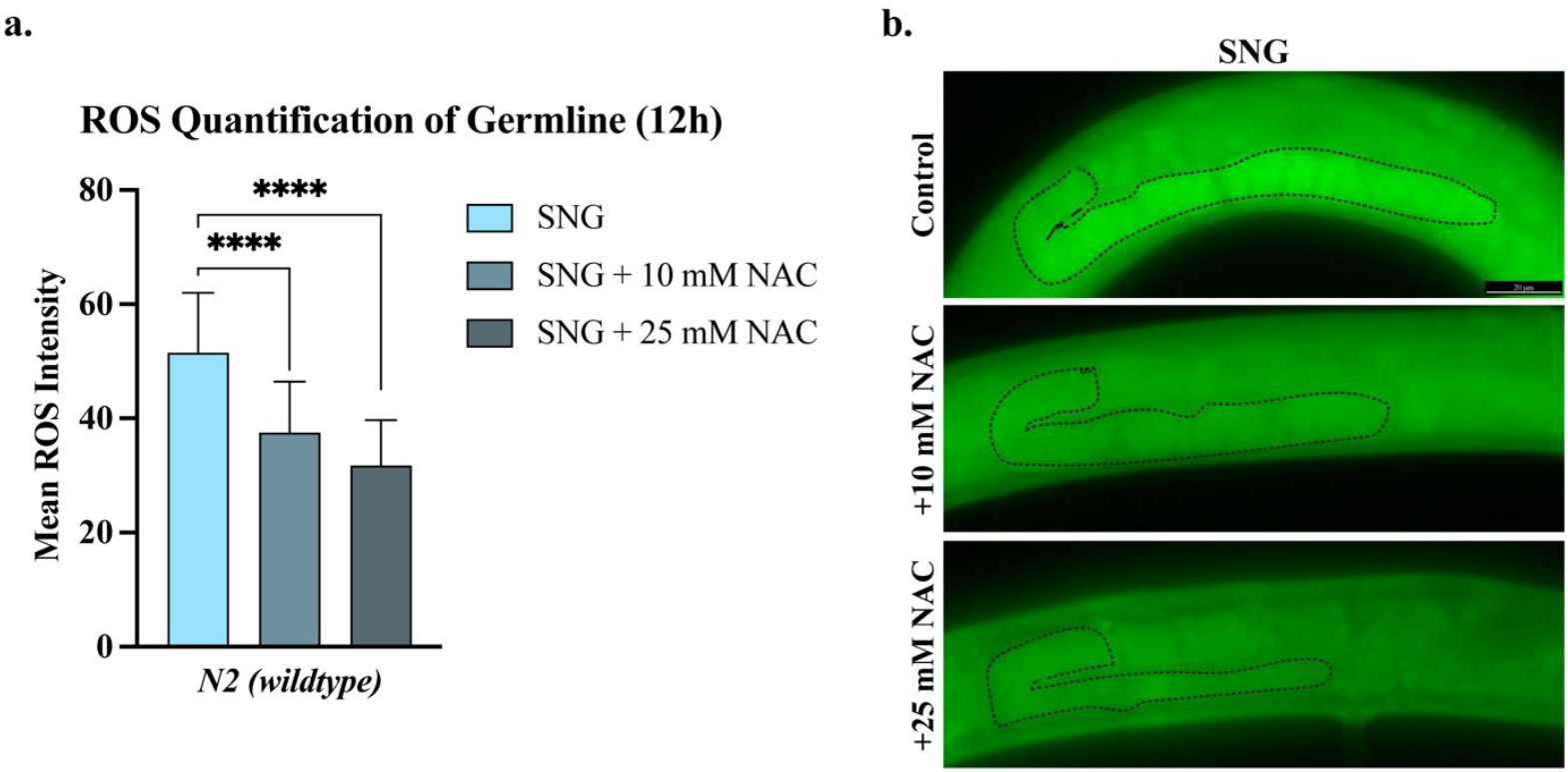
Treatment with the ROS scavenger NAC decreased germline ROS levels in SNG-treated worms. **a**. Treatment with both 10 mM and 25 mM NAC significantly decreased germline ROS levels (**** p<0.0001) in sanguinarine-treated worm **b**. Germlines of SNG-treated worms stained with CellROX Green, without and with 10 mM or 25 mM NAC.

We next asked whether attenuation of oxidative stress by NAC affected the apoptotic response. Quantification of apoptotic corpses in *ced-1::gfp* worms revealed that NAC treatment also led to a significant reduction in SNG-induced germline apoptosis compared to SNG-treated controls (p<0.0001; **Figure 6**). Together, these results demonstrate that oxidative stress is a critical mediator of SNG-induced germline apoptosis.

**Figure 6.**
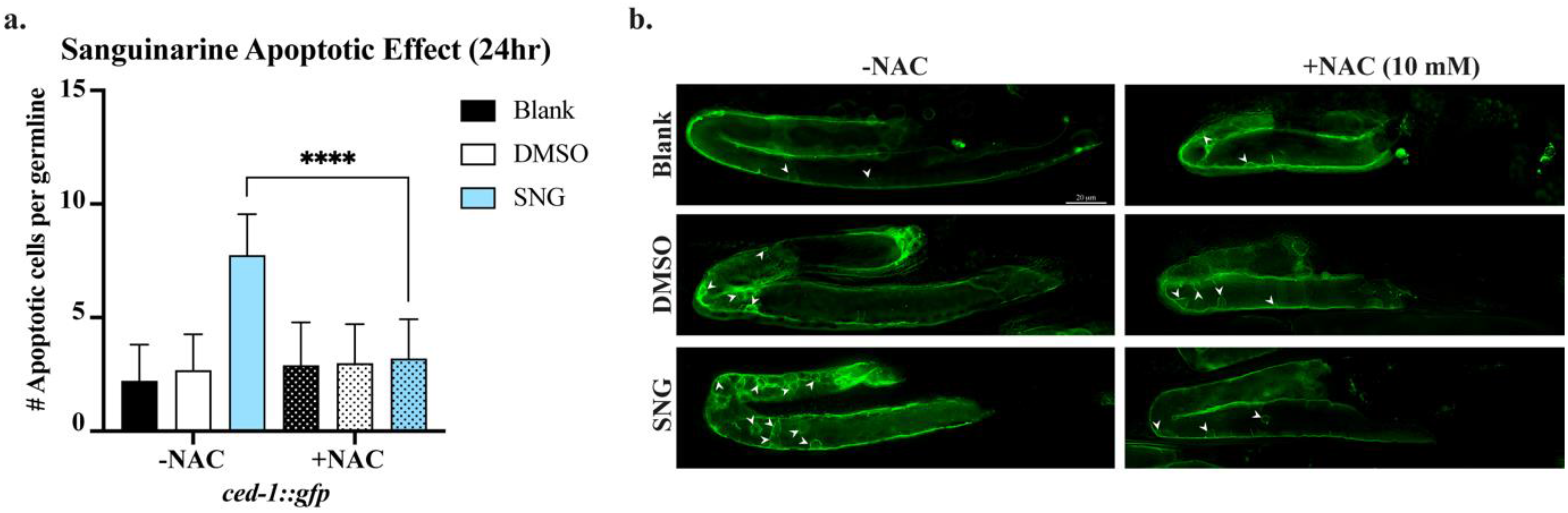
Treatment with ROS scavenger NAC suppresses apoptotic phenotype in SNG-treated worms. **a**. A significant reduction in germline apoptosis upon 10 mM NAC cotreatment with SNG (**** p<0.0001). **b**. Microscopic images of apoptotic cells in the germlines of *ced-1::gfp* worms in response to Blank, DMSO, and SNG treatments with and without the addition of 10 mM NAC. Arrowheads mark apoptotic cells.

## Discussion

In this study, we identify sanguinarine as a potent inducer of germline apoptosis through activation of the conserved DNA damage response in *C. elegans*. Using genetic and cytological approaches, we show that SNG activates the canonical ATM/ATR–CHK-1 checkpoint pathway, leading to CEP-1/p53–dependent, caspase-mediated apoptosis in the germline. Notably, SNG-induced apoptosis is highly tissue-specific and is not related to developmental apoptosis in somatic tissues, as our analysis on developmental apoptosis revealed no increase in apoptotic corpses in the developing larvae. We further demonstrate that SNG treatment results in a pronounced accumulation of reactive oxygen species selectively within the germline, and that pharmacological scavenging of ROS suppresses both oxidative stress and germline apoptosis. Together, these findings reveal that sanguinarine engages a coordinated DNA damage checkpoint and oxidative stress response to selectively eliminate compromised germ cells *in vivo*.

Mechanistically, our data place SNG-induced germline apoptosis downstream of activation of the canonical DNA damage checkpoint signaling pathway mediated by the ATM/ATR kinases axis. We show that SNG treatment induces phosphorylation of ATM/ATR substrates, as detected using an antibody that specifically recognizes the conserved phospho-(S/T)Q motif. Consistently, SNG treatment also leads to phosphorylation and activation of the conserved DNA damage checkpoint kinase 1, CHK-1, a key downstream effector of ATM/ATR signaling.

Importantly, our data reveals a striking and unexpected feature of SNG exposure in *C. elegans*, a robust increase in ROS. In contrast to mammalian cells, where SNG induces widespread oxidative stress across multiple cell types, the oxidative response in *C. elegans* is highly restricted to the proliferative germline, as somatic tissues remain largely unaffected. This observation challenges the general view that SNG-induced oxidative stress represents a basic toxicity effect of the compound and instead highlights a strong cellular-context dependency. It is worth noting that in adult *C. elegans*, the germline is the only mitotically active tissue, suggesting that SNG-induced ROS accumulation occurs in proliferative cells. Thus, the oxidative stress imposed by SNG appears to be linked not simply to compound exposure per se, but to features specific to dividing cells, which may be related to their metabolic state, redox balance, or replication process. Importantly, pharmacological inhibition of S-phase progression failed to suppress SNG-induced ROS production. Treatment with 5-fluorouracil (5-FU), which effectively blocks DNA replication and germ cell proliferation, did not abolish ROS induction upon SNG exposure. These findings indicate that SNG-induced oxidative stress is not a direct consequence of DNA replication itself but rather reflects a broader metabolic or signaling state associated with proliferation.

Consistent with a causal role for oxidative stress, the SNG-induced increase in ROS was completely abolished upon treatment with the ROS scavenger *N*-acetylcysteine. Elevated oxidative stress is well known to promote genomic instability and to activate the ATM/ATR-mediated DNA damage checkpoint, ultimately inducing p53-dependent programmed cell death. Notably, scavenging ROS not only suppressed ROS accumulation but also fully rescued the apoptotic phenotype induced by SNG exposure. Collectively, these findings place oxidative stress as a critical mediator linking SNG treatment to DNA damage–induced apoptosis.

All in all, our data reveal a mechanistic framework for sanguinarine-induced germline apoptosis *in vivo*, in which SNG exposure imposes oxidative stress, activates DNA damage checkpoint signaling, leading to p53-dependent cell death specifically within the germline of *C. elegans*.

## Materials and Methods

### *C. elegans* strains and maintenance

*C. elegans* strains were maintained on Nematode Growth Media (NGM) plates seeded with *E. coli OP50* bacteria as previously described [44]. All strains were grown at 20 °C unless otherwise stated. The *C. elegans* strains used in this study: Bristol N2 (wildtype), MD701: *bcIs39 [lim-7p::ced-1::GFP + lin-15(+)]*, EPD077: *cep-1(Ig12501);[lim-7p::ced-1::GFP + lin-15(+)]*, KX84: *ced-3(n2452) IV; bcIs39 V*, and CB3203: *ced-1(e1735)I*.

### Sanguinarine treatment

Synchronized larval stage 1 (L1) worms were grown until they reached the larval stage 4 (L4) and then were treated with various sanguinarine doses ranging from 8 to 25 μM for 24 hours in liquid culture. The initial sanguinarine dosages tested were 8 µM, 10 µM, 15 µM, 20 µM, and 25 µM, diluted from a 10 mM sanguinarine stock dissolved in 1 M dimethyl sulfoxide (DMSO). Following the dosage response, 25 μM sanguinarine was chosen and used to carry out subsequent experiments. A volume of DMSO equal to the highest used drug concentration was used as a control; the final DMSO concentration was 2.5 mM.

### Apoptosis scoring

Apoptotic cells were measured as previously described [37, 40, 42, 45]. In brief, treated worms were mounted on slides with a 3% agarose pad and immobilized with 50% levamisole. Apoptotic corpses were monitored and scored in the germline using the Leica 3D-Thunder Imager Fluorescent microscope equipped with differential interference contrast (DIC) optics. Apoptotic cells were detected and counted per germline using both DIC optics and the expression of CED-1::GFP using the GFP filter.

### Developmental apoptosis assay

The engulfment-defective *ced-1*(*e1735*) strain was used to assess developmental apoptosis. Synchronized L4 staged worms were placed on NGM plates containing dissolved DMSO or 25 μM sanguinarine, supplied with OP50 bacteria, and incubated at 20 °C. After 24 hours of treatment, the adult worms were washed off their respective treatment plates with M9 and bleached to extract embryos. The following day, hatched L1 worms were mounted on slides containing agar pads with 50% levamisole. Apoptotic cells in the pharyngeal region of the L1-staged worm were counted for each condition using DIC microscopy [37].

### Immunostaining

Synchronized L4-staged N2 worms were treated with DMSO, Hydroxyurea, Sanguinarine, or left untreated (Blank) for 24 hours, followed by germline extraction and immune staining as described previously [37, 40, 42, 45-47]. The following primary antibodies have been used with 1:200 dilution: Anti-Phospho-CDK-1 (Thr14, Tyr15) Mouse Monoclonal Antibody (MABE229, EMD Millipore), Anti-Phospho-CHK-1 (Ser345) Rabbit Monoclonal Antibody (S.48.4) (MA5-15145, Thermo Scientific), and Anti-Phospho-ATM/ATR Substrate motif ((S/T)Q) (6966, Cell Signaling Technology). The secondary antibodies used in this study were: Donkey Anti-Mouse IgG H&L (Alexa Fluor® 488) (ab150105, Abcam) (1:200), Donkey Anti-Rabbit IgG H&L (Alexa Fluor® 568) (ab175470, Abcam) (1:200). Additionally, a DAPI stain (1:1000) was used for DNA staining.

### ROS detection and quantification

To measure ROS levels generated upon sanguinarine treatment, the CellROX™ Green Reagent (Invitrogen, CAT #C10444) was used. N2 wild-type worms at the L4 stage were treated in liquid culture with various conditions for 12 or 24 hours. Worms were then washed three times with M9 buffer, and the samples were fixed in 2% paraformaldehyde dissolved in M9 on a rotator platform at 300 rpm for 30 minutes at RT. Fixed worms were then washed three times with PBS and stained with 5 mM CellROX™ Green solution for one hour at RT. After staining, worms were washed three times with PBS and then mounted on agar pads for imaging. Images were captured with Thunder Imager using a 20X objective and the GFP filter. ImageJ was used to quantify ROS levels after SNG treatment. The GFP intensity of the regions of interest (ROIs) was measured by outlining the ROIs and calculating their mean intensity using the Lasso tool. For germline ROS measurement, the GFP and DIC images were superimposed to determine the germline structure, and the Mean Gray Value was measured from the GFP panel.

### NAC treatment

For ROS quantification upon ROS scavenger treatment, N2 worms at the late L3 stage were pretreated with 10 or 25 mM NAC for 12 hours in liquid culture. Once worms reached the L4 stage, DMSO or SNG was added and incubated for 12h, after which germline ROS levels were measured as described in the “ROS detection and quantification” section. For apoptosis scoring, *ced-1::gfp* worms at the L4 stage were co-treated with 10 mM NAC and 25 μM SNG, DMSO, or blank for 24 hours, then apoptotic cells were quantified using fluorescent microscopy as described.

### 5-Fluorouracil treatment

Wildtype N2 worms were treated for 4 hours in 50 mM 5-fluorouracil in liquid culture, then their respective drugs (Blank, DMSO, Sanguinarine) were added for an additional 12 hours. The ROS levels were measured using CellROX™ Green as described.

### Statistical analysis

Statistical analysis and graph generation were performed using GraphPad Prism version 10.1.0 (GraphPad Software, San Diego, CA, USA). The means of the data were compared between the control and treatment groups, and significance was assessed using one-way ANOVA. The obtained p-value was compared with the p-value threshold set: *p < 0.05 – Statistically significant; **p < 0.01 – Highly statistically significant; ***p < 0.001 – Very highly statistically significant; and ****p < 0.0001 – Extremely statistically significant.

## Acknowledgments

We would like to thank all the members of the Pourkarimi Lab for their constructive comments and feedback throughout this project. We would also like to acknowledge the *Caenorhabditis Genetics Center* (CGC), funded by the NIH (National Institute of Health) Office of Research Infrastructure Programs (P40OD010440), for providing *C. elegans* strains. This work and the Pourkarimi Lab are funded by the Qatar Foundation and the College of Health and Life Sciences at Hamad Bin Khalifa University (HBKU).

## Author Contributions

Conceptualization, investigation, methodology, visualization, writing of the original draft, R.E.; conceptualization, investigation, M.I.; conceptualization, investigation, Z.A.; conceptualization, investigation, T.A.A.; validation, review, S.U.; conceptualization, project administration, resources, supervision, validation, writing, review, and editing of the original manuscript, E.P. All authors have read and agreed to the published version of the manuscript.

## Figures

**Supplemental Figure S1.**
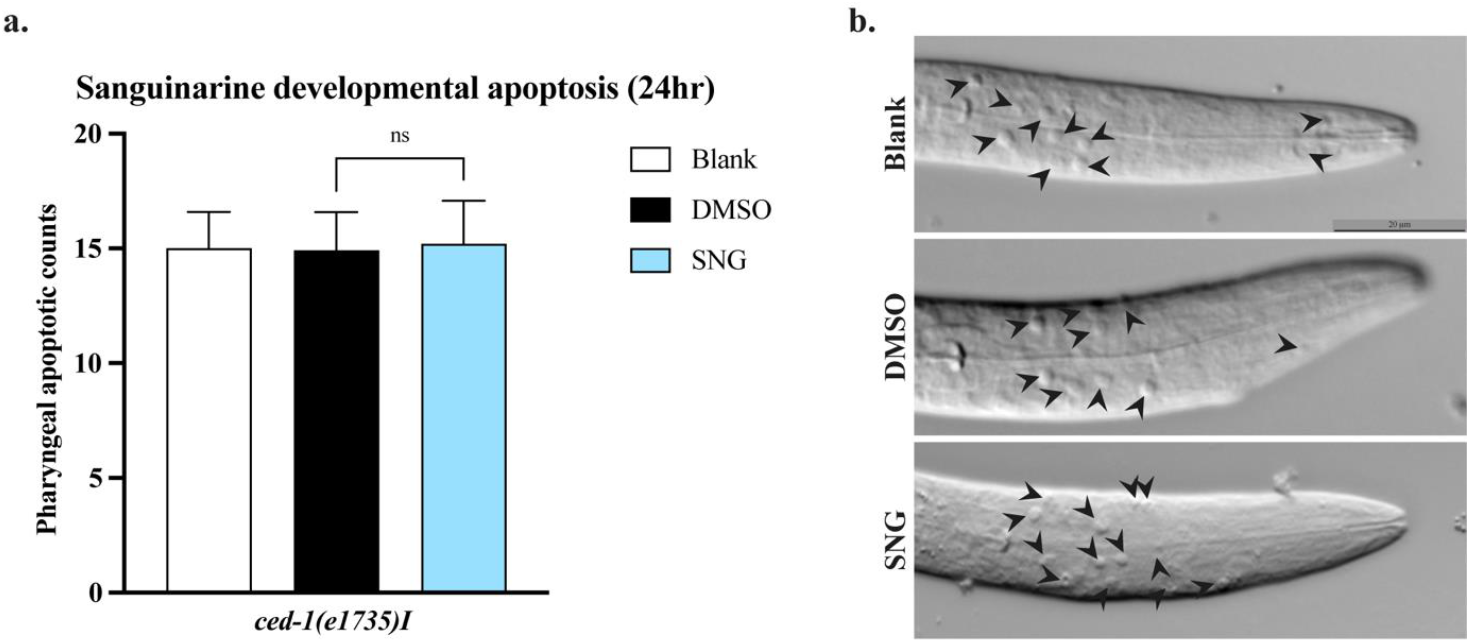
Developmental apoptosis in the progenies of SNG-treated worms. **a**. Quantification of pharyngeal apoptotic cells in L1 larvae showing no significant difference between DMSO- and SNG-treated groups. **b**. Representative images of apoptotic cells in the pharyngeal region of L1-staged *ced-1(e1735)* worms following Blank, DMSO, or SNG treatment.

**Supplemental Figure S2.**
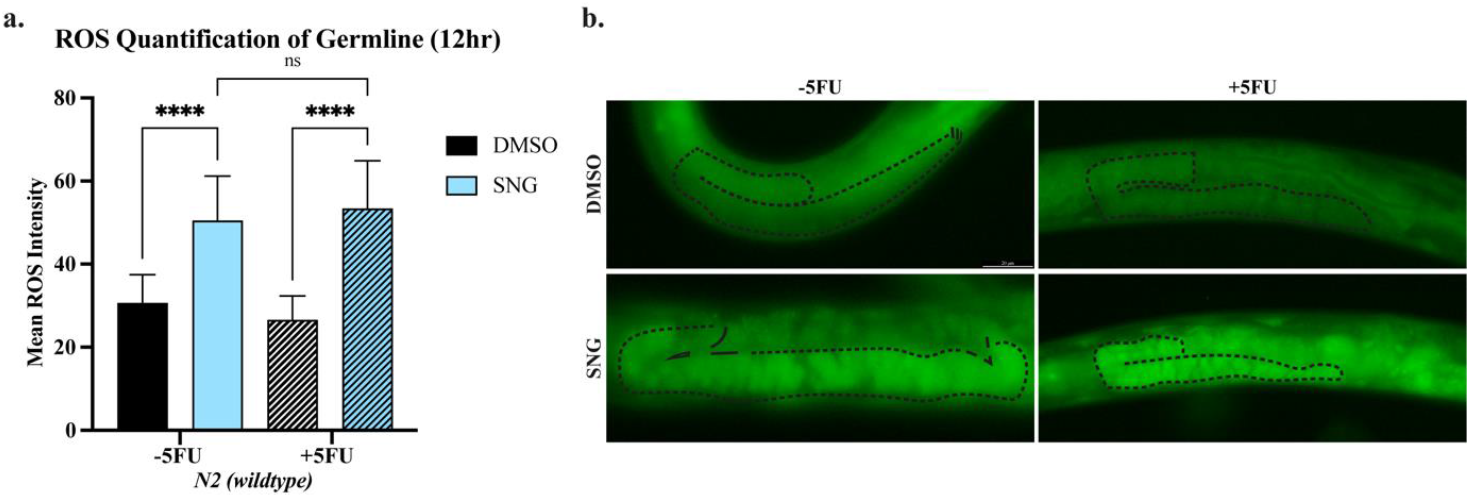
Blocking germline mitosis does not alter SNG-induced ROS levels. **a**. Quantification of ROS levels showing no significant difference between SNG-treated worms in the presence or absence of 5-fluorouracil (5FU). **b**. Representative images of CellROX Green–stained germlines following 12 h of SNG treatment. Worms in the right panel were pretreated with 5FU to inhibit germline mitosis prior to SNG exposure.

